# Defining the design requirements for an assistive powered hand exoskeleton

**DOI:** 10.1101/492124

**Authors:** Quinn A. Boser, Michael R. Dawson, Jonathon S. Schofield, Gwen Dziwenko, Jacqueline S. Hebert

## Abstract

The goal of this study was to identify design criteria for the development of an assistive powered hand exoskeleton by consulting with potential end users. Structured interviews with clinicians and patients with hand impairment were carried out and the results were tabulated. Three participants with impaired hand function also underwent a quantitative measurement session regarding hand function. The objective of the measurement sessions was to understand the characteristics, abilities and limitations of the upper limb of individuals who could benefit from a hand exoskeleton device, in order to better define design criteria and control options for such a device. For the most part, clinicians and participants with hand impairment agreed on expectations for a hand exoskeleton device on topics including important grasp patterns, wear time, and grip strength. However, their expectation seemed to diverge on the topic of control, where clinicians felt simple reliable control strategies would be preferred, but patients desired intuitive control. This research has identified key features of hand exoskeleton design requirements that will need to be met in order to have acceptable clinical translation to patient populations. Including endusers in the design of such a device is essential for successful patient-oriented technology development.

## 1. Introduction

The human hand is an intricate machine that is vital to interact with our environment. The importance of dextrous hand function becomes most evident when impairment of hand function occurs. Limited hand function can significantly impede the ability of an individual to perform activities of daily living and dramatically impact quality of life. In Canada, it is estimated that 62,000 incidences of stroke occur yearly [1], with nearly 80% suffering partial upper limb paralysis [2]. Furthermore, over 37,000 Canadians are living with compromised upper limb function resulting from spinal cord injury [3]. Beyond these populations, numerous neurological disorders and traumatic injuries can result in compromised hand function.

Powered hand exoskeletons are a newly emerging technology that have demonstrated promising results in alleviating functional challenges associated with hand impairment or weakness [4,5]. These systems attach to segments of the hand, and actively assist digit flexion and extension to aid in performance of functional grasping tasks by applying forces to appropriate areas of the user’s digits. Therefore, they can help the user restore movement by guiding their digits to specific hand positions or grasping patterns; movement that would typically be challenging or not possible for these individuals to independently achieve and maintain stable grasp.

Several hand exoskeleton designs have been described in scientific literature [6,7,8,9,10]. However, the majority have not been validated in patient populations or translated beyond a laboratory environment. Those that have are high cost and typically operate in a purely therapeutic capacity, often being tethered to a computer displaying a virtual reality environment. To date, only one commercially available assistive device exists that allows the user to be untethered and able to be used for day-to-day tasks [11]. To our knowledge, it is not yet freely available for purchase outside of the United States. Consequently, the functional benefits of hand exoskeleton technology remain largely inaccessible to clinicians and patients. Furthermore, designs for exoskeletons are quite varied and it is not clearly described which populations would benefit most from them. Ideally, a clinically accessible assistive hand exoskeleton device would come from a set of well-defined design specifications derived from the input of end users, specifically patients and the clinical professionals who work with them.

## 2. Study Objectives

The purpose of this study was to define objective design requirements for an assistive powered hand exoskeleton capable of clinical or long term use. This aim was pursued by focused interviews with individuals with hand impairment (potential end users of the device), and clinicians at the Glenrose Rehabilitation Hospital who work with such patients. Further design requirement criteria were gathered by characterizing hand function of three participants with hand impairments.

## 3. Methods

### 3.1. Recruitment

**Clinician Participants:** Clinician participants were recruited through the Glenrose Rehabilitation Hospital through the lead occupational therapist on this project. Potential participants were approached in person and recruitment was done through ‘word-of-mouth’, with an attempt to engage clinical staff with experience working with patients with hand impairment.

**Patient Participants:** Patient participants were recruited from patient roles at the Glenrose Rehabilitation Hospital. The clinical coordinators and occupational therapist at the hospital were asked to identify potential participants that would be capable of participating in focus groups and/or individual interviews, based on their cognitive and language ability, who met the criteria of impaired hand function. Subject recruitment was on the basis of a convenience sample, without specific matching. There is no reason to believe that the clinical benefits of an assistive hand exoskeleton would be affected by gender, age or race. Therefore, participants recruited were of any race or gender. Inclusion criteria were adults (18 to 75 years of age) with non-progressive or static hand impairment, including diagnoses such as brain injury, spinal cord injury, and nerve injury (total n=8). Exclusion criteria were cognitive impairments or language barriers that would inhibit their ability to comprehend and respond to the interview questions.

All participants provided signed informed consent and this study was approved by the University of Alberta Health Research Ethics Board. One participant consented by video conference and his signed consent form was mailed in to the researchers.

### 3.2. Design Criteria Interview Sessions

Interviews were conducted with small groups (< 3 participants), or with individual participants when it was not possible to attend a small group. Interview questions were divided into two sections: 1) design criteria questions, which were aimed at understanding and quantifying specific design expectations and requirements, and 2) open discussion questions, which were intended to help the researchers appreciate the participants’ views on advanced assistive technology, and visions of how a hand exoskeleton might be implemented and used. Two of the design criteria questions involved demonstrations; one made use of a video showing different possible hand open/close speeds, and for the other, different servo motors were run to demonstrate the amount of noise that they generate. The specific questions used in the interviews are provided in Appendix A.

Present at each interview were: one main researcher (the interviewer); the interviewee(s); and one or two other lab members, to assist with recording and transcribing the interview.

#### 3.2.1. Clinician Interview

One group interview was completed with 6 clinicians who were employed with The Glenrose Rehabilitation Hospital or Alberta Health Services at the time of the interview. The specific questions used in the clinician interview are provided in Appendix A.1. The group consisted of occupational therapists (OT) and hand rehabilitation specialists who work with patients with impaired hand function. Six clinicians participated in the interviews. Their years of experience ranged from 5-15 years working in occupational therapy or hand therapy, and included specialization in one or more of the following patient populations: populations with traumatic injuries (including spinal cord and peripheral nerve injury), brain injuries, and stroke.

#### 3.2.2. Interviews with Hand Impaired Participants

A total of 7 interviews were completed with 8 participants with impaired hand function on at least one side of their body. The specific questions used in the interviews are provided in Appendix A.2. Eight participants were recruited from the following diagnostic categories: stroke, spinal cord injury (SCI), and brachial plexus injury. When possible, we attempted to schedule participants from the same diagnostic category together, in order to complete the interviews in small groups. However, scheduling conflicts often did not allow for this. The breakdown of interviews was as follows:

– Interview 1: one participant with SCI, completed via video call
– Interview 2: two participants with brachial plexus injury
– Interview 3: one participant with SCI
– Interview 4: one participant with SCI
– Interview 5: one participant with brachial plexus injury
– Interview 6: one participant who had suffered a stroke
– Interview 7: one participant who had suffered a stroke

### 3.3. Hand Characterization Measurement Sessions

Separate measurement sessions were conducted with three participants with impaired hand function:

– Par05 - 67-year-old male with a spinal cord injury (SCI) three years prior, affecting roots C4 through C7 (assumed C5 spinal cord level of injury with shoulder spared, limited elbow function on right arm). Measurements were taken on right (weaker) side, because he indicated that, that was the hand he would prefer to use a hand exoskeleton with, if it were available.
– Par06 - 62-year-old male with a brachial plexus injury three years prior, affecting right arm
– Par07 - 27-year-old male who had experienced a stroke, seven years prior, affecting right side

The objective of these sessions was to define characteristics of the participants’ hands which would be relevant to hand exoskeleton design criteria, as well as assess the viability of different mechanisms for potentially triggering a hand exoskeleton device. The full protocol for these sessions is included in Appendix B, and is summarized below.

#### 3.3.1. Assessment of Hand Function

The participants were first asked to attempt to hold their impaired hand in a number of different grasp patterns and perform different movements in order to assess their current abilities and which grasp patterns they would need the most help achieving. An experimenter recorded whether the participant was able to obtain each grasp pattern or movement and photographs were taken. It was also recorded whether the experimenters were able to move the participants thumb into opposition, if they were not able to actively do it themselves.

#### 3.3.2. Range of Motion

Active and passive range of motion was assessed for flexion of the index finger using a finger goniometer. Range of motion was first assessed on the index finger, if the other fingers had a similar movement pattern then range of motion was not measured for them in the interest of time. Also note that passive hyper-extension was not measured, as a hand exoskeleton would likely not need to hyperextend the fingers.

Active wrist range of motion was also measured, to assess the viability of a flexion sensor at the wrist as a trigger for activating a hand exoskeleton device.

#### 3.3.3. Grip Force

Participants’ unassisted grip force was measured in cylinder (diameter = 70 mm), tip -to-tip, tripod and lateral key grip patterns. Participants were asked to grasp a load cell with different attachments for the different grip patterns. The experimenters would sometimes help the participant to shape their hand around the load cell and attachments; however, they would not help them to apply force. Participants were asked to squeeze the load cell as hard as they could and then relax (not maintain the force). Three repetitions were completed when possible.

#### 3.3.4. External Force Required to Flex/Extend Index Finger

A load cell attachment was designed to allow the experimenters to pull the participants’ index finger into extension, and push it into flexion, while measuring the force required to do so. Participants were asked to relax and not resist the movement or actively assist with it. Movement was stopped if it became uncomfortable for the participant, or the experimenter felt uncomfortable with the level of resistance. Three movements were examined: index finger extension (of all three joints simultaneously), flexion of the MCP joint, and flexion of the PIP and DIP joints while the MCP joint was stabilized. For one participant thumb opposition was also assessed. At least three repetitions were completed for each movement.

#### 3.3.5. Forearm Electromyography (EMG)

Surface EMG readings were recorded from sites on the participants’ forearm flexor and extensor muscles. The participants were asked to complete a number of movements including wrist flexion and extension, making a fist, extending all fingers, and pointing their index finger. The participants were told that if they were not able to actively complete the movements, it was okay, and to still imagine doing them, and to express the intention of the movement.

For analysis of the EMG data in this report, the mean absolute value of the data was calculated using a moving window and plotted, a representative contraction was selected for each movement, and an average reading was estimated for the corresponding agonist and antagonist muscles. Noise level was estimated before the onset of the signal.

## 4. Results

### 4.1. Design Criteria Interview Sessions

#### 4.1.1. Envisioned Use

##### 4.1.1.1. Activities & Grasp Patterns

Clinicians emphasized the importance of being able to get the thumb into opposition during this section of the interview. They also agreed that you could probably have movement of digits 2/3 and 4/5 linked to move together.

The participants with hand impairment listed a wide variety of activities that they would like to perform with the help of a hand exoskeleton. The activities that were mentioned are summarized in Figure 1, as well as the number of participants who mentioned each activity. The importance of thumb opposition was also brought up by one participant with hand impairment.

**Figure 1:**
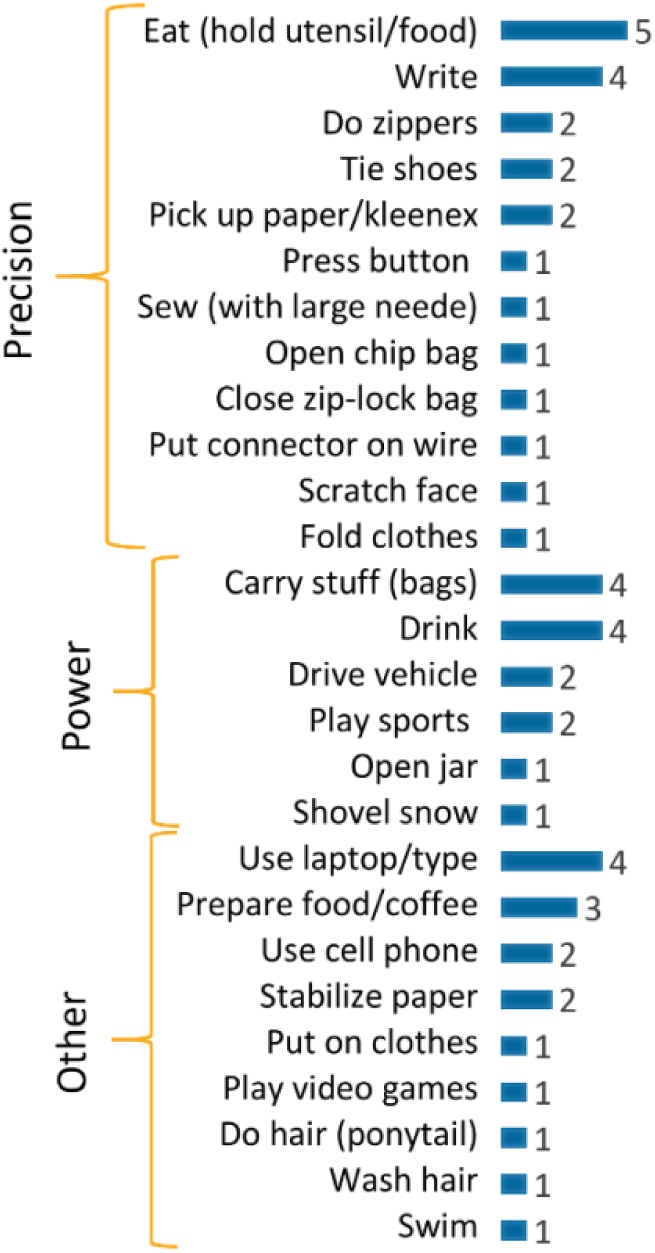
Activites that participants with hand impairment indicated they would like to be able to do with the assistance of a powered hand exoskeleton device. Numbers on the left indicate the number of participants (out of 8) who mentioned each activity.

##### 4.1.1.2. Grasp Strength

This question elicited a wide variety of responses. Some participants with hand impairment were only envisioning lifting light objects such as utensils, while one participant was imagining heavier objects (up to 1 kg). A theme that came up in with both clinicians and participants with hand impairment was being able to lift the weight of a drink (average coffee, beer, 750 ml drink).

##### 4.1.1.3. Usage Time

The clinicians felt that battery life for a hand exoskeleton device should be at least 6-8 hours. Four of eight participants with hand impairment said they would like to wear the device “all day” or “constantly,” if possible. Two participants said that they would wear the device for part of the day ("5-6 hours", and “afternoon"). Two participants said the amount that they would wear it would depend on how comfortable or bulky it was. Importantly, participants did not envision this as a device that they would want to put on and take off multiple time throughout the day, but one that they would keep for an extended period of time. This expectation comes with some other implied requirements that came up in the interviews (in addition to battery requirements):

– Ideally, the hand exoskeleton would be waterproof, or the design would be such that individuals could easily wash their hands without taking the device off completely
– The hand exoskeleton device should be slight enough that an individual can put on or take off a jacket while wearing it (this was mentioned as being especially relevant in the Canadian climate)

##### 4.1.1.4. Range of Motion

Most participants with hand impairment indicated that they would want to be able to close their hand enough to perform pinch grip (or one mentioned carrying a bag). With respect to opening, participants mentioned being able to open hand wide enough to grasp a coffee cup or reach for something above them on a shelf.

##### 4.1.1.5. Closure/Opening Speed

Most participants with hand impairment said they would be satisfied with 2 second opening/closing speed. A couple participants indicated that they would prefer 1 second. An interesting challenge was noted here as even participants who had a stroke indicated that they would prefer faster closure speeds, but anecdotally, their spasticity and tone might not allow it, as the current movement capable with their hand was of slow speed.

##### 4.1.1.6. Control

Clinicians had some concerns about using EMG to trigger a hand exoskeleton device, particularly the risk of unwanted movements, and being difficult to learn. Clinicians seemed to agree that it would be best if the mechanism for triggering the device was modular such that it could easily connect it to a button for example, in addition to EMG. We did not specifically ask participants with hand impairment about control strategy; however, a couple of patient participants did ask about it, and one mentioned that they would not use the device if they had to push a button to control it.

##### 4.1.1.7. Donning/Doffing

We did not specifically ask participants about donning/doffing of a hand exoskeleton device. However, during the open discussion portion of the interviews, in response to the question “What do you think would be some of the most important factors for you to want and to use an exoskeleton during your day?” 6 of the 8 participants with hand impairment indicated that being able to put on and take off the device independently would be important to them.

#### 4.1.2. Physical Properties

##### 4.1.2.1. Weight and Bulk

Questions on size elicited a range of responses. Most participants with hand impairment (7/8) had some size that they would consider too bulky to wear (the other participant said they would be okay with the device being bulky now, because they would expect it to get smaller as technology improves). However, the size that individuals thought was acceptable varied. The most extreme example was one participant who imagined a hand exoskeleton having the bulk of a smaller ‘leather glove, with some extra bulk at the wrist.’ Most participants agreed that a 5×5×3 cm (3 cm = height) block was too bulky for the back of the hand.

Most participants with hand impairment (7/8) said 200 g would be manageable on the hand (one participant said 200 g might still be too heavy). A couple of participants thought 500 g would be manageable.

Most participants with hand impairment indicated that they would be okay with moving bulk from the hand onto the forearm wherever possible (and would prefer it). Two participants indicated that they would still like to be able to cover their arms with sleeves (either for aesthetic purposes, or weather protection). A couple of participants said they would be willing to try a design where some weight was located on the body (ex: on a belt), but others were strongly against this.

##### 4.1.2.2. Noise

A couple participants indicated that they would not care about noise, and would wear a hand exoskeleton device which was noisy, as long as it was helping them. However, most participants with hand impairment thought at least one of the motors demonstrated was too noisy to use in public.

### 4.2. Hand Characterization Measurement Sessions

#### 4.2.1. Assessment of Hand Function

Table 1 provides a summary indicating which grasp patterns and hand movements the participants were able to complete. Par05 (SCI) was not able to complete any functional grasp patterns or wrist movements. The experimenters were partially able to passively pinch their thumb and index finger together; however, there was too much muscle stiffness to completely bring the thumb into opposition against the index and middle finger. Par06 (brachial plexus injury) was able to partially or completely complete all grasp patterns and movements except wrist extension against gravity, but struggled to obtain the precision grip patterns (tip-to-tip and tripod). Par07’s (stroke) hand was clutched in a fist due to spasticity; however, their fingers would curl around a cylinder or sphere if they were first passively opened to accept the object.

**Table 1:** Summary of which grasp patterns and movements participants were able to perform with their impaired hand. ✓ indicates ‘yes’, ✘ indicates ‘no’, and ✓ indicates ‘partially’

|  | Par05<br>(SCI) | Par06<br>(brachial<br>plexus) | Par07<br>(stroke) |
| --- | --- | --- | --- |
| Active hand opening | ✗ | ✓ | ✗ |
| Active grip - cylinder | ✗ | ✓ | ✓<sup>2</sup> |
| - hook | ✗ | ✓ | ✗ |
| - spherical | ✗ | ✓ | ✓<sup>3</sup> |
| - tip-to-tip | ✗ | ✓<sup>1</sup> | ✗ |
| - tripod | ✗ | ✓<sup>1</sup> | ✗ |
| - lateral key | ✗ | ✓ | ✗ |
| Active wrist extension, gravity eliminated | ✗ | ✓ | ✗ |
| Active wrist extension against gravity | ✗ | ✗ | ✗ |
| Active wrist flexion, gravity eliminated | ✗ | ✓ | ✗ |
| Active wrist flexion against gravity | ✗ | ✓ | ✗ |
| Hand palm down - fingers extended | ✗ | ✓ | ✓<sup>4</sup> |
| **Passive thumb opposition** | ✓ | ✓ | ✓ |

<sup>1</sup> Required a lot of concentration
<sup>2</sup> Hand in full fist at rest, but not able to actively open to take cylinder shape
<sup>3</sup> Fingers can curl around ball when placed in hand, but thumb not in opposition
<sup>4</sup> With help from experimenter

#### 4.2.2. Range of Motion

Active and passive range of motion in flexion for the three participants is summarized in Table 2.

**Table 2:** Summary of participant active and passive range of motion in flexion

| Joint |  | Par05 (SCI) |  | Par06 (brachial plexus) |  | Par07 (stroke) |  |
| --- | --- | --- | --- | --- | --- | --- | --- |
|  |  | Active | Passive | Active | Passive | Active | Passive |
| Wrist |  | - <sup>1</sup> |  | -31°- 46° |  | - <sup>1</sup> |  |
| Index | MCP | - <sup>2</sup> | 0°- 65° | 5°- 80° | 0°- 88° | - <sup>3</sup> | - <sup>4</sup> |
|  | PIP | - <sup>2</sup> | 16°- 54° | 36°- 100° | 0°- 94° | - <sup>3</sup> | - <sup>4</sup> |
|  | DIP | - <sup>2</sup> | 5°- 49° | 20°- full fist | 0°- full fist | - <sup>3</sup> | - <sup>4</sup> |
| Middle | MCP | * | 0°- 45° | * | * | * | * |
|  | PIP | * | 15°- * | * | * | * | * |
|  | DIP | * | 18°- * | * | * | * | * |
| Ring | MCP | * | 0°- 34° | * | * | * | * |
|  | PIP | * | 26°- * | * | * | * | * |
|  | DIP | * | 15°- * | * | * | * | * |
| Pinky | MCP | * | 0°- 36° | *- 34° | *- 42° | * | * |
|  | PIP | * | 36°- * | *- 100° | *- 105° | * | * |
|  | DIP | * | 8°- * | *- 80° | * | * | * |

\* For the middle, ring and pinky finger, if movement limit was similar to the index, joint angles were not recorded in the interest of time
<sup>1</sup> Not recorded because participant did not have active wrist movement
<sup>2</sup> No active movement, resting joint angles were MCP ≈ 30°, PIP ≈ 25°, DIP ≈ 15°
<sup>3</sup> Very limited active movement, hand fully closed in fist when in resting position
<sup>4</sup> Not recorded – Passive extension was difficult to measure, but experimenters could get hand mostly open if it was moved very slowly. Hand could be fully closed.

Par05 (SCI) had almost no active range of motion and some muscle stiffness that resisted passive motion. Their hand could not be fully opened or closed passively. Par06 (brachial plexus injury) could not fully extend their fingers actively, but could almost make a full fist, except for the pinky finger. Passively their hand could be fully opened and almost fully closed, except for the pinky finger. Par07’s hand (stroke) was closed in a fist even in resting position. They were not able to actively extend their fingers. The experimenters were able to obtain passive extension but it was very difficult to measure the joint angles as the movement had to be very slow, and opening force had to be maintained or the fingers would flex again.

Of the three participants, only Par06 (brachial plexus injury) was able to obtain active wrist flexion/extension, making them the only one of the three participants that would be able to use a flex sensor at the wrist as a triggering mechanism for a hand exoskeleton device.

#### 4.2.3. Grip Force

Force data collected from the grip force measurements is shown in Figure 2 and Figure 3 for Par06 and Par07 respectively. Three repetitions of force application were completed for each grasp pattern whenever possible and are shown in the figures.

**Figure 2:**
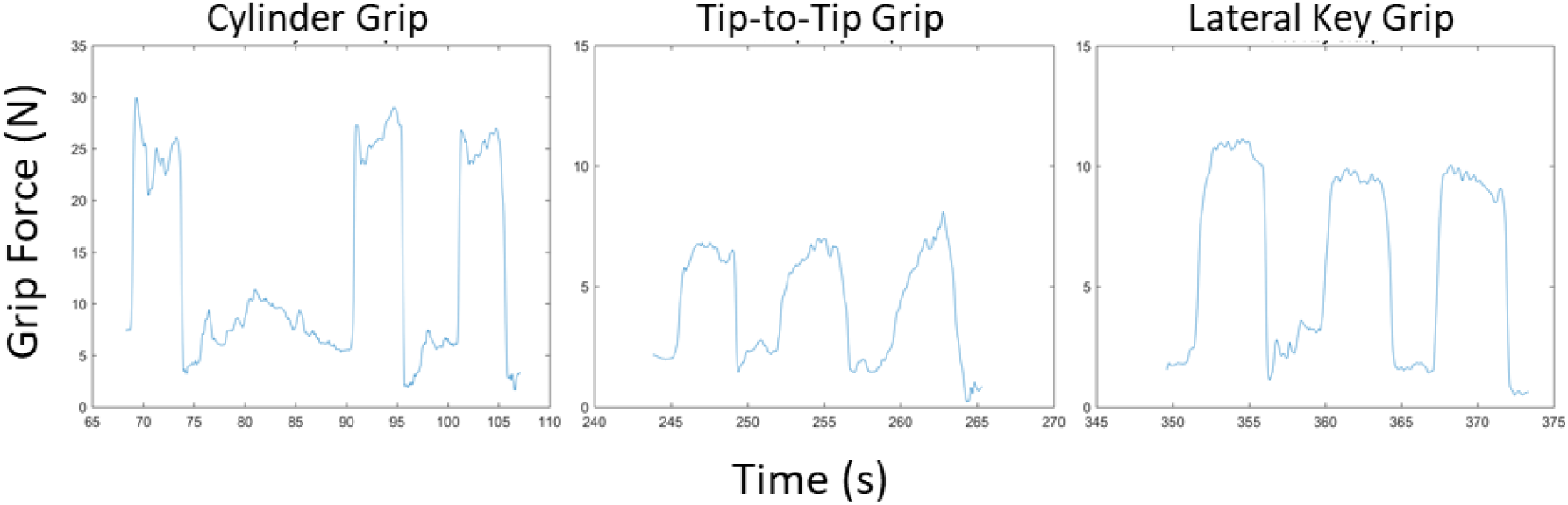
Grip force data for Par06 (brachial plexus injury)

**Figure 3:**
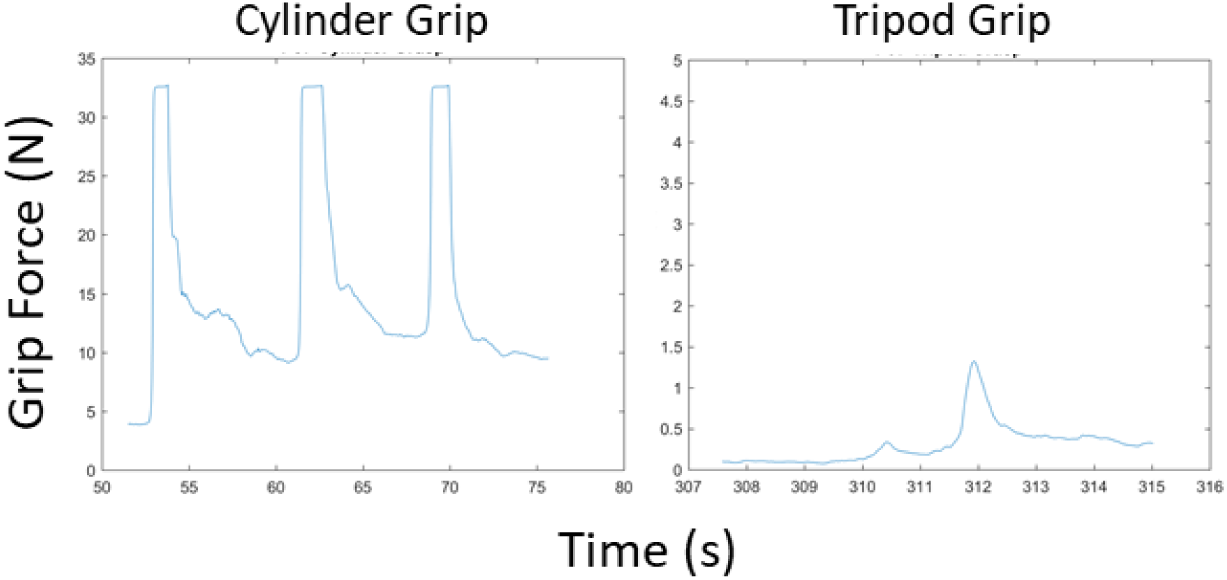
Grip force data for Par07 (stroke)

Par05 (SCI) was not able to apply grip force in any of the grip patterns. Par06 (brachial plexus injury) was able to complete the cylinder, tip-to-tip and lateral key grip, but not the tripod grip. For tip-to-tip, the experimenters helped to position the load cell with attachments in the participant’s hand, but they were able to squeeze it without assistance. Par07 (stroke) was able to complete the cylinder grip and one repetition of the tripod grip with help from the experimenters to position the measurement device in their hand for both. Maximum grip forces, averaged across the repetitions, are summarized in Table 3.

**Table 3:**
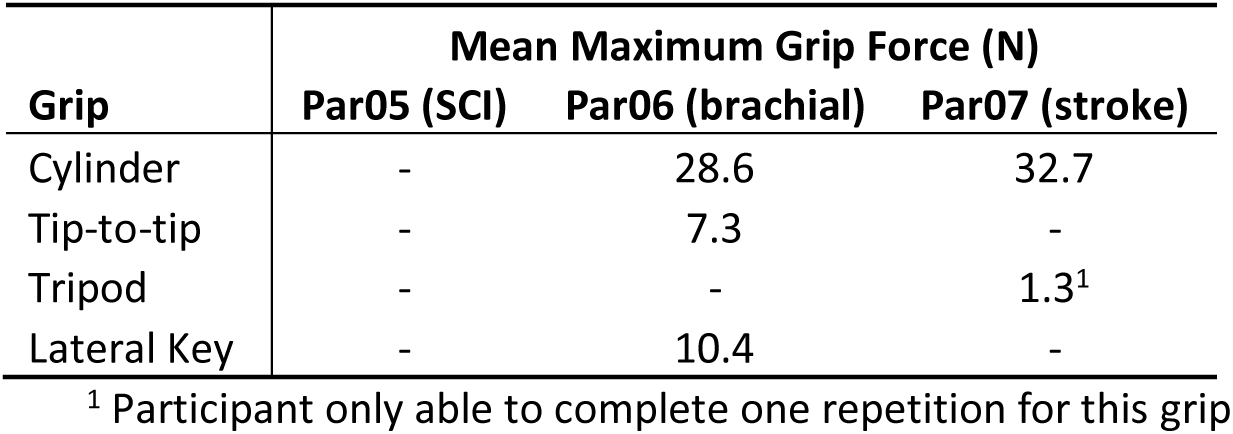
Summary of participant maximum grip force, averaged across three repititions when possible

#### 4.2.4. Force Required to Flex/Extend Index Finger

Plot of the applied force required to flex and extend the index finger vs. time are provided for each participant in Figure 4. In most cases three repetitions of the movement were repeated. For Par07 (stroke) finger extension, five repetitions were repeated because of perceived variability in muscle spasticity. Force required to push the thumb into opposition was also measured for Par06 (brachial plexus injury), because this participant emphasized the desire to have a device which could actively move their thumb into opposition.

**Figure 4:**
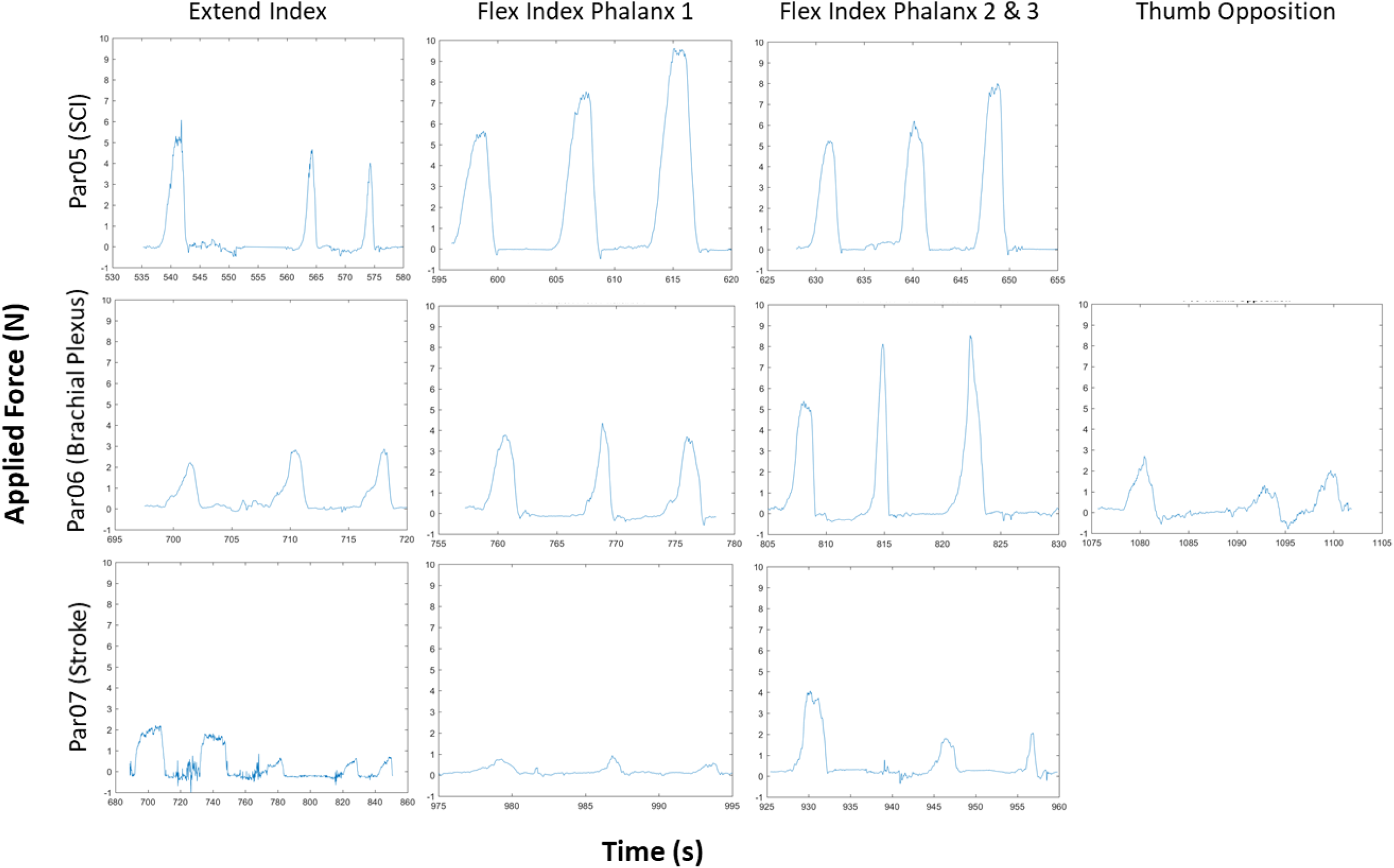
Plots of applied force required to flex and extend index finger for each participant, as well as force required to move thumb into opposition for Par06. Rows of plots are associated with the same participant, while columns of plots are associated with the same movement

**Figure 5:**
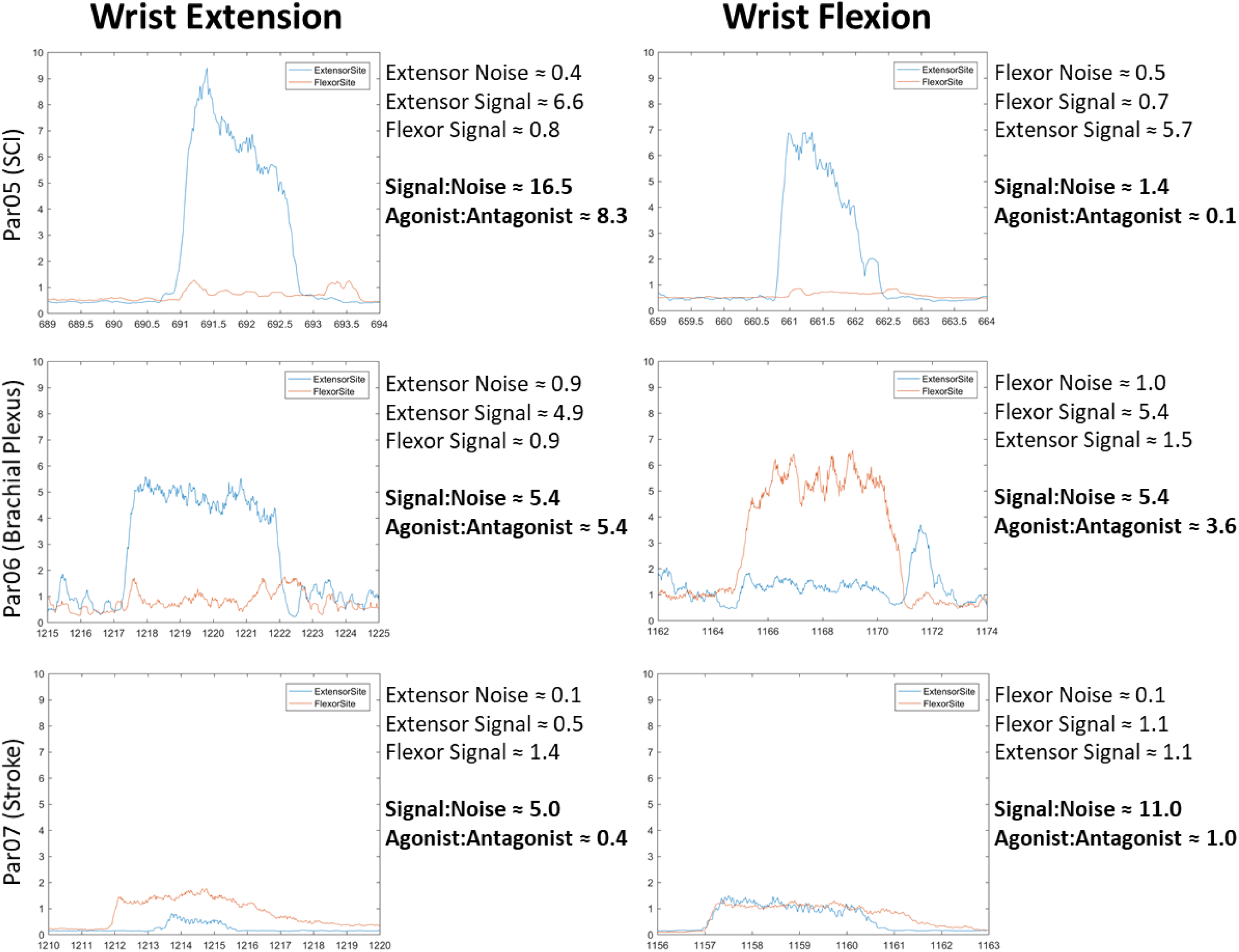
Surface EMG signals recorded from participants’ forearms during wrist flexion and extension. Figures show mean absolute value of the raw EMG signals, calculated using a moving window. Rows of plots are associated with the same participant, while columns of plots are associated with the same movement.

The peak applied force and overall movement duration (time from onset of increasing force, to force returning to level of noise) was extracted for each movement repetition. Average peak force (over the 3 or 5 repetitions), absolute maximum peak force, and average movement duration are summarized in Table 4.

**Table 4:**
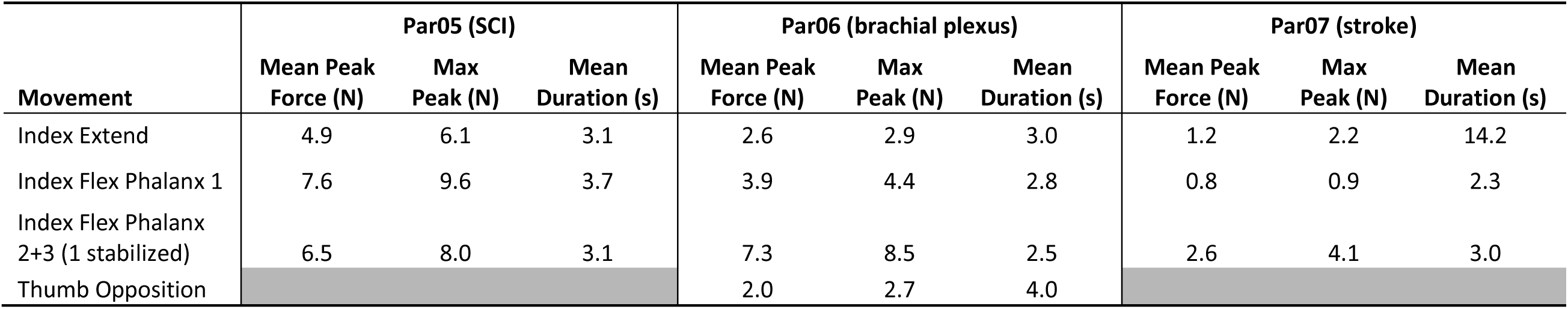
Summary of mean peak force and maximum peak force required to flex/extend index finger for each participant and oppose thumb for Par06, as well as mean movement durationcs

| Movement | Par05 (SCI) |  |  | Par06 (brachial plexus) |  |  | Par07 (stroke) |  |  |
| --- | --- | --- | --- | --- | --- | --- | --- | --- | --- |
|  | Mean Peak Force (N) | Max Peak (N) | Mean Duration (s) | Mean Peak Force (N) | Max Peak (N) | Mean Duration (s) | Mean Peak Force (N) | Max Peak (N) | Mean Duration (s) |
| Index Extend | 4.9 | 6.1 | 3.1 | 2.6 | 2.9 | 3.0 | 1.2 | 2.2 | 14.2 |
| Index Flex Phalanx 1 | 7.6 | 9.6 | 3.7 | 3.9 | 4.4 | 2.8 | 0.8 | 0.9 | 2.3 |
| Index Flex Phalanx 2+3 (1 stabilized) | 6.5 | 8.0 | 3.1 | 7.3 | 8.5 | 2.5 | 2.6 | 4.1 | 3.0 |
| Thumb Opposition |  |  |  | 2.0 | 2.7 | 4.0 |  |  |  |

For Par05 (SCI) the greatest forces were required to flex phalanx 1 of the index finger, with an overall peak force of 9.6 N. From visual inspection of Figure 4, it appears as if the force required to flex the index increases with each movement repetition.

For Par06 (brachial plexus injury) the greatest forces were required to flex phalanxes 2 and 3 of the index finger, with an overall peak force of 8.5 N for this movement. However, this participant was able to actively perform this movement (see hook grip in Table 1), and so a hand exoskeleton might not be required to provide this full force. This participant was more concerned with having a device which could assist them with thumb opposition, which required an average peak force of 2 N, and an overall maximum of 2.7 N.

For Par07 (stroke) required forces were relatively low compared to the other participants (with a maximum of 4.1 N required to flex phalanx 2 and 3 of the index, with phalanx 1 stabilized). The main thing to note for this participant was the much greater amount of time required for finger extension. Based on results from the interview sessions, the experimenters aimed for a 2 second movement time. However, movement times ranged from 1.4 to 5 seconds for all movements except Par07 finger extension. For Par07, muscle spasticity made it impossible to extend the fingers at this speed, and finger extension could only be achieved if the experimenters moved much more slowly (average movement duration = 14.2 s). The time required to extend the index finger appeared to decrease with each successive repetition, based on the 5 trials that were collected.

#### 4.2.5. Forearm Electromyography (EMG)

Of the movements examined, wrist flexion and extension consistently produced the most promising EMG signals for all participants, consequently these were the only movements analyzed for this report. The participants completed three repetitions of wrist flexion as well as extension. Mean absolute value of the raw EMG signals from a representative trial for each participant are shown in Figure 4, along with hand average estimates of signal and noise values, and corresponding signal-to-noise and antagonist-to-agonist ratios. Note that all gains were set to 100 for Par05, and 40 for Par06 and Par07.

It can be observed that Par05 was able to produce a strong extensor signal. However, when asked to perform wrist flexion, they also activated their wrist extensor muscles. This implies that they could use one-site EMG control to trigger a hand exoskeleton, but would not be appropriate for conventional two-site control.

Par06 was the best candidate for conventional two-site EMG control, as we were able to obtain strong extensor and flexor signals with a signal-to-noise ratio of approximately 5.4 for each.

Par07 had difficulty isolating flexion/extension signals, and instead tended to co-contract their forearm muscles, more so during wrist flexion. This pattern could be used to trigger a device using one-site control, or possibly some sort of pattern recognition algorithm, but would not be useful for conventional two-site EMG control.

## 5. Discussion

The objective of this study was to identify design criteria for the development of a powered hand exoskeleton by consulting with potential end users of such a device - patients with hand impairment, and clinicians who work with such patients. For the most part, clinicians and participants with hand impairment agreed on expectations for a hand exoskeleton device on topics including important grasp patterns, wear time, and grip strength. However, their expectation seemed to diverge on the topic of control – clinicians recognized the challenges associated with more complicated control strategies such as EMG, and suggested starting with simpler strategies like a button. When the topic came up with participants with hand impairment, some of them expressed that push button control would turn them off of using the device, and expected more intuitive control options. Both precision and power grips were emphasized as important, with a variety of weight requirements; at least the weight of an average drink seemed to be a key strength requirement. Wear time was requested to be 6-8 hours with ideally continuous use, not requiring frequent donning and doffing, and therefore able to be washed for hygiene and able to wear clothing over it. Acceptable bulk and weight ranged from a small glove to less than a 3 cm block extending on the back of the hand.

The three participants that underwent the detailed hand measurements all had quite different hand characteristics, suggesting that there would not be a single ideal hand exoskeleton that would fit all participants with hand impairment. Par05 (SCI) would benefit most from a device which was focused on applying forces to flex the fingers, and had stronger motors that would need to be able to exert forces up to 10 N to overcome muscle stiffness. This would probably not be achievable with a fully wearable device. Because this participant used a wheelchair, it might be possible to mount a larger battery to the wheelchair. However, this would make such a device unusable for ambulatory individuals. Par06 (brachial plexus injury) would benefit most from a device focused on active control of thumb opposition. They would also be a good candidate for a larger number of control options, particular using proximal muscle signals for activation. Par07 (stroke) would benefit from a device which focused on extension of the fingers at a very slow speed, and controlled flexion and thumb opposition positioning.

It is possible that there would be more similarities with a larger participant pool. However, based on the interview sessions, there was considerable variability within and across populations. This emphasizes that hand function and hand characteristics are based on a number of factors including severity, level and time since injury, as well as presence of spasticity, contracture, or flexibility of the impaired hand. When designing hand exoskeletons for the purpose of providing an assistive device, design features will need to consider target population(s), and the variability that will exist between individuals with the same diagnosis.

### 5.1. Future Work and Recommendations

This research has identified key features of hand exoskeleton design requirements that will need to be met in order to have acceptable clinical translation to patient populations. Including end-users in the design of such a device is essential in patient oriented technology development. In addition to identifying design criteria, this research identified several prototypes of hand exoskeletons that were in various stages of development (Appendix C). We anticipate that the many existing hand exoskeleton devices which are currently under development will benefit from the end user feedback which has been compiled in this study.

Other authors have reported limited aspects of design requirements, for example Smaby et al, 2004 [12], documented that for a set of functional activities “pinch force requirements ranged from 1.4 N to push a button on a remote to 31.4 N to insert a plug into an outlet". Of the tasks studied by Smaby et al, “9 of 12 required less than 10.5 N". Therefore 10 N of force may be a reasonable goal to aim for in a hand exoskeleton. However, the results of our study emphasized that consideration needs to be given to the force needed to initiate movement of the digits and overcome resistance due to muscle stiffness or spasticity (which was up to 10 N in the participants we studied), in addition to the force required for a functional grip.

In addition, Hume et al, 1990 [13] identified functional requirements for finger range of motion during a number of practical activities. The activities examined by Hume et al were fairly consistent with the activities requested by the participants in our study (Figure 1), and so the results of this study could likely be applied to hand exoskeleton design, in that to be function a powered hand exoskeleton device would not need to be able to fully flex the fingers. The maximum flexion angles observed by Hume et al during their functional tasks were 73°, 86° and 61° for the MCP, PIP an DIP joints respectively, and the average functional angles were 61°, 60° and 39°. However, our study also highlighted that some individuals with impaired hand function may not have access to this functional range of motion, even in passive motion, due to muscle stiffness or contracture (Table 2). This fact must also be considered in hand exoskeleton design so that these devices do not injure their wearers by trying to move their fingers past their passive range of motion.

To our knowledge, hand open/close speed required for a hand prosthesis or assistive device to be considered functional has not previously been quantified. Participants in our study mostly selected 2 seconds as an acceptable amount of time for a hand to spend opening/closing. However, when we assessed the force required to extend the index finger of one of our participants who had experienced a stroke, we found that the movement had to be much slower to not activate the participants’ spasticity. Ideally a successful hand exoskeleton device might have an adjustable speed to account for this.

Overall, based on the results of this design requirement study, it seems that if powered hand exoskeletons are going to succeed as clinical assistive devices, there may need to be a variety of different successful devices targeting different populations with specific characteristics, or it may be that a custom modular approach needs to be pursued in order to allow individualization of design.

A current scan of the literature and commercial products has identified a few hand exoskeleton devices with potential to be applied as functional assistive devices (Appendix C). With the advance of soft robotics and smaller powered devices, it is possible that some of the user requirements may be met in existing devices or iterations in the near future.

In particular, the Myomo MyoPro^®^ is a powered elbow/wrist/hand orthosis, which is available in the United States, and is marketed as an assistive device for individuals with “brachial plexus injury, brain or spinal cord injury, CVA stroke, multiple sclerosis or amyotrophic lateral sclerosis.” While the website presents compelling anecdotal information and preliminary [14] and preliminary clinical evidence with stroke patients [15], clinical evidence that this device is useful for all of the targeted patient groups is lacking. Additionally, research labs such as The Seol National University BioRobotics Laboratory [4,10,16] and Rehabilitation Engineering Lab out of Zurich [17] have also presented some potentially promising hand exoskeleton designs. However, many of these designs have only been evaluated with healthy individuals or a very limited number of individuals with impaired hand function (1 or 2 – See Appendix C). Indeed, clinical trials demonstrating the efficacy of these devices in the target patient populations are needed in order for this technology to move forward to iterations that will be acceptable to patient populations and clinicians, in a manner that improves daily function.

## Acknowledgments

We would like to thank the Glenrose Rehabilitation Hospital clinicians, patients, and research department for their support of this project. Quinn Boser was financially supported by a TD Bank Financial Group Health Sciences Interdisciplinary Research Studentship award.

## Appendix A. Interview Questions

### A.1 Clinician Interview Questions

Design Criteria Questions:

1. How large do you feel this item can be on the hand? Should we aim to “minimize size” or do we want to settle on a hard number?

The researchers will bring a measuring tape, digital calipers, example electronics enclosures, example actuators, and example battery packs to provide context to the interviewees.
2. Are there foreseeable issues with placing componentry proximal, perhaps on the forearm?
3. How heavy to we think is too heavy to be functional? Again, are there foreseeable issues with moving componentry more proximal?

– A range of weighs from ~100g to 100kg will be brought to the meeting for the interviewee group to manipulate and add context to this criterion.
4. How do you foresee the user using this device? What types of hand grasping patterns to you predict as being the most beneficial to the user?

– Here the concept of digit actuation and the number of actuators required to achieve various grasp patterns will be discussed.
5. What range of motion do you see as being functional? Do you feel the ability to achieve a fully closed fist is necessary, or can it be a “loose fist,” or partially closed hand? What role do we see the thumb play? Should it be splinted, manually moved or actively moved with a motor?

– Members of the interviewee group may be asked to demonstrate their view of a functional range of motion. The researchers will bring a goniometer and measuring tape to fully quantify any demonstrated movements.
6. What types of objects do you predict your clients may want to grasp and hold? How heavy do you predict these objects may be?

– The concept of balancing grip strength and actuator size/power requirements will be highlighted here.
– A range of weighs from ~100g to 100kg will be brought to the meeting for the interviewee group to manipulate and add context to this criterion.
7. What are your thoughts on battery/device runtime? How many hours of the day might you predict a patient may use an exoskeleton device?
8. We have a number of ways we can trigger such a device to open and close (such as push buttons, EMG sensors, flex sensors and motion sensors). Do you have any particular thoughts on how a client may want to activate their device? Can you identify any obvious limitations?

– Here the researchers should also address the idea of modularity in which different triggering mechanisms may be applied for different patients.
9. How adjustable or modular would you like an exoskeleton device to be? Do you prefer one size fits all, or is some customization acceptable? If so to what degree?
10. In terms of technical specifications (mechanical, electrical or otherwise) do you feel that we have overlooked anything? If so please let us know and explain the significance.

Open Discussion:

1. Have you used advanced rehabilitation technology before (automated or computerized assistive or rehabilitative technologies)?

1. If so which technologies? What were your impressions and why? Can you elaborate on its strengths and weakness and why they are strengths and weaknesses?
2. If not, why? What barriers have prevented you from accessing such technologies?
2. How would you foresee yourself and your patients interacting with a hand exoskeleton system in the clinic or at home?
3. What specific requirements do you believe are necessary to allow this technology to be successful in the clinic and at home?
4. Today we spoke a lot about hand exoskeletons, what about this technology seems most promising to you? What about it make you the most apprehensive?
5. If we were to brain storm some outcome measures to evaluate the success of this device, what types of activities would we perform? What types of tests would you like to see? What do you think these test may show?
6. Do you have any additional comments that you would like to express?

### A.2 Patient Interview Questions

Design Criteria Questions:

1. If a hand exoskeleton was available, how would you like to use it? What types of activities would you like to perform? What types of hand grasping patterns would you most likely use?
2. If you had a hand exoskeleton, what types of objects would you want to grasp and hold? How heavy of an object would you like to be able to grab and hold?

– A range of weighs from ~100g to 1kg will be brought to the meeting for the patient to manipulate (if they are capable) and add context to this criterion. If not useful data can still be gained from the description of objects they wish to be able to manipulate.
3. If you were able to use an exoskeleton, how long each day do you think you would like to use it?

– Here the researchers should also address the idea of modularity in which different triggering mechanisms may be applied for different patients.
4. How wide would you like to be able to open your hand and how far would you like to be able to close your hand?

– If the patient has a healthy hand, they will be asked to demonstrate, if not a research will demonstrate for them. The researchers will bring a goniometer and measuring tape to fully quantify any demonstrated movements.
5. We have prepared a demonstration of different hand closure speeds. Would you be able to watch these videos of a hand closing and tell us which closing speeds seem acceptable, and which would be too slow?

– The researchers will bring a video of a hand closing at a few different speeds (0.5 seconds, 1 second, 2 seconds…). The patient’s impressions of the closing speeds will be recorded
6. How large do you feel this item can be on the hand? Should we aim to “minimize size” or do we want to settle on a hard number?

– The researchers will bring a measuring tape, digital calipers, example electronics enclosures, example actuators, and example battery packs to provide context to the interviewees.
7. Do you think you would have any issues with placing parts of the device on your forearm or closer to your body?
8. How heavy to we think is too heavy? If we moved some of the weight to your forearm or closer to your body do you think you would have any issues?

– A range of weighs from ~100g to 1kg will be brought to the meeting for the patient to manipulate and add context to this criterion.
9. We have brought a number of small motors with us today. Would you be able to listen to them and tell us which you think make an acceptable amount of noise if they were to be used in a hand exoskeleton that you wore daily?

– The researchers will bring a number of actuators and demonstrate them to the patient. The patient’s impressions of the noise they make will be recorded.

Open Discussion:

1. What do you think would be some of the most important factors for you to want and to use an exoskeleton during your day?
2. In your opinion, what would make this device a “success” for you? What things what functionality or options would make you want to use this device or not want to use it?
3. Today we spoke a lot about hand exoskeletons, what about this technology seems most promising to you? What about it makes you the most doubtful?
4. Have you used advanced rehabilitation technology before (automated or computerized assistive or rehabilitative technologies)?

a. If so which technologies? What were your impressions and why? Can you elaborate on its strengths and weakness and why they are strengths and weaknesses?
b. If not, why? What barriers have prevented you from accessing such technologies?
5. How would you foresee yourself using a hand exoskeleton system? Would you wear it all day or just sometimes?
6. Do you have any additional comments that you would like to express?

## Appendix B. Hand Measurement Session Protocol

<u>Introduction:</u>

– Review consent form
– Give overview of session (first we will take pictures and measurements, then ask them to do some grasping, then put sensors on the skin)
– Ask if sensitive to adhesives or medical tape
– Give them a time estimate, ask if they want a break at any point

<u>Hand Photographs:</u>

1. Place white photo backdrop down on table (or surface that participant can reach)
2. Place L-shaped goniometer/ruler down
3. Ask participant to place forearm and hand (up to elbow if they can) on surface and take photos in following configurations (ask them to attempt – we just want to capture what they are currently capable of doing):

- Side of hand (ulnar side down)

Resting
Voluntary open - fingers extended as far as they are able
Voluntary close - cylinder – “Can you make a fist”

– hook - “Can you curl your fingers, like this?"
– spherical (ask them to close hand around actual sphere)
– tip-to-tip - “make a circle /w tip of index & thumb"
– tripod
– lateral key - “pinch thumb against the side of finger" * with last two grasp patterns; note on data form whether they can get thumb into opposition, if not, note whether it can be passively moved into opposition
– Wrist flexion
– Wrist extension
- Hand palm down

– Fingers extended (do **not** apply passive over pressure)
– Wrist extension – looking for radial deviation
- Hand palm up

– Wrist flexion – looking for ulnar deviation

<u>Hand Geometry:</u>

**If** participant is able to hold their hand flat on a surface (palm down)

1. Carefully trace their hand – keeping the pencil normal to the page and as tight against their hand as possible
2. Make a marking at the locations of interphalangeal joints as well as ulnar and radial styloid processes
3. PIP to MCP of each finger
4. Measure 5^th^ MCP to wrist (mountable space)

**Otherwise**

1. Use soft tape measure to take relevant measurements to complete hand geometry section of data form

<u>Active & Passive Range of Motion:</u>

Wrist:

1. Measure range of active wrist flexion and extension using goniometer, and record on data form

Finger Extension:

**If** participant’s fingers can be passively extended so that hand is flat

→ Record this fact, no need to take measurements

**Otherwise**

→ For each finger that cannot be fully extended passively, measure the limits of passive extension for the MCP, PIP and DIP using finger goniometer, and record on the data form

Finger Flexion:

1. Ask the participant to place their hand on the table (or a surface they can reach), on its side (ulnar side down) in a resting position

*If it is comfortable, goal is to measure with wrist at normal resting position (~20° extension), if not comfortable, measure at whatever wrist angle is comfortable
2. Ask the participant to curl their fingers into a fist as much as they are able
3. For the index finger: measure the limit of active flexion for the MCP, PIP and DIP

→ After measuring each joint, push gently on the distal segment to measure the range of passive flexion - “Now I am going to gently push to see how much farther we can move this joint. Let me know right away if there is any pain or discomfort"
4. **If** all other fingers move with the index, note this, and only measure active and passive ranges for each MCP. **If not**, all joints for each finger that does not move with the others.

<u>Grip force</u>:

1. Begin logging data on data logging computer
2. Make sure grip force attachments are secured to load cell and load cell is sending data to the computers
3. Tare the load cell
4. Ask the participant to grip the load cell using each of the following grasp patterns

- Cylinder grip
5. Increment the label number
6. Change the attachments on the load cell to the ones for fine grasp patterns
7. Increment label *again* number to signal start of new grasp
8. Ask the participant to grip the load cell using each of the following patterns (remember increment the label number at the end of each grasp, and *again* when they are ready to start next grasp)

- Tripod
- Tip-to-tip
- Lateral key
9. *Record the current label number for each grasp pattern on the data form

<u>Force required to open/close hand:</u>

1. Switch the load cell attachment to the finger puller
2. Ask participant to rest their hand on table, ulnar side down
3. Ask the participant to hold their hand in a resting position with fingers slightly closed (as much as they are able, while still being relaxed)
4. Gently stabilize their metacarpal with your hand
5. Place the finger puller under the third phalanx of their index finger, close to the DIP joint
6. Increment label number to indicate start of pulling
7. Gently pull the finger into extension, as much as you are able without causing them discomfort (aim for 1 second opening time)
8. Change label number to indicate end of motion, repeat 3 times
9. Ask the participant to hold their hand in a resting position with fingers slightly extended (as much as they are able, while still being relaxed)
10. Gently stabilize their metacarpal with your hand
11. Place the finger pusher side of the attachment on the back of the **first** phalanx of their index finger
12. Increment the label number to indicate the start of pushing
13. Gently push the first phalanx into flexion, as much as you are able without causing any discomfort (aim for 1 second closing time)
14. Change label number to indicate end of motion, repeat 3 times
15. Gently stabilize the first phalanx of their index finger
16. Place the finger pusher side of the attachement on the back of the **third** phalanx of their index finger
17. Increment the label number to indicate the start of pushing
18. Gently push the second and third phalanx into flexion, as much as you are able without causing any discomfort (aim for 1 second closing time)
19. Change label number to indicate end of motion, repeat 3 times
20. Repeat process with any one fingers that seem like they might be interesting or give different values

<u>Electromyography data</u>:

1. Ask the participant to try flexing and extending their wrist
2. Look for muscle bellies
3. Place electrodes around the forearm, on muscle bellies
4. Increment the label number
5. Ask the participant to perform the following movements (3 repetitions each):

- Wrist flexion
- Wrist extension
- All finger flexion
- All finger extension
- Index finger flexion
- Index finger extension
- Finger abduction
6. Change the label number in between each movement
7. Try to take notes on any electrodes that seem to be getting a good signal during specific movements

## Existing Hand Exoskeleton Designs

### Literature Summary

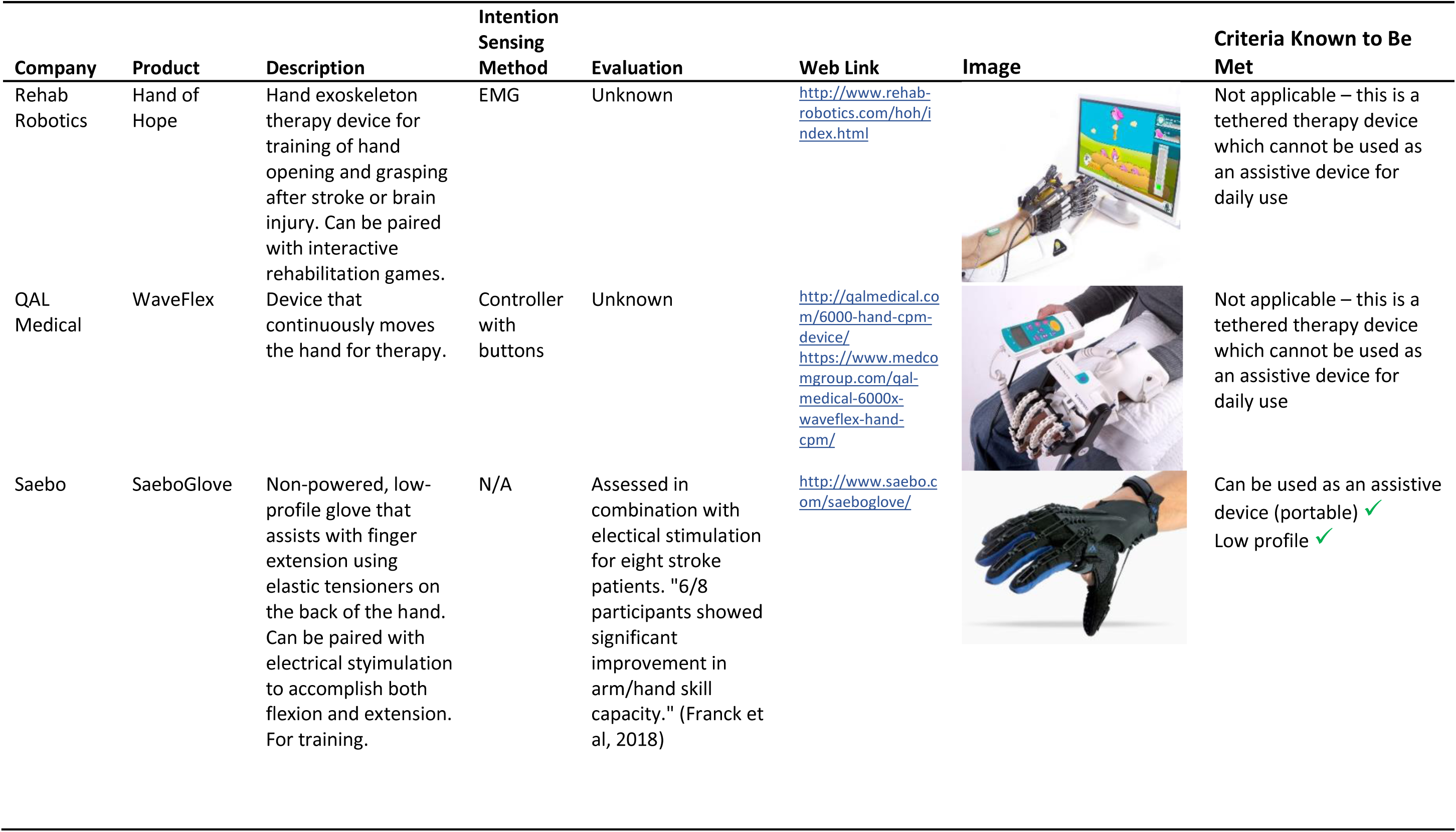

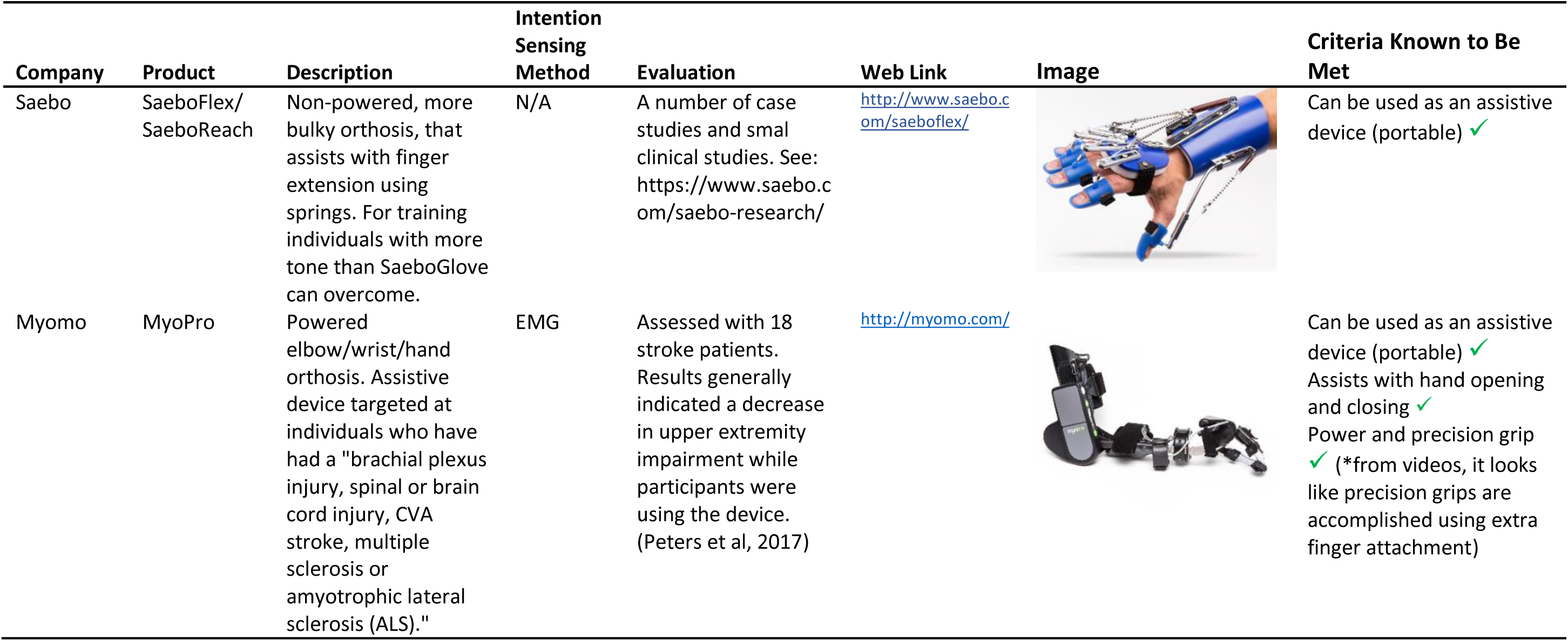

### Literature Summary

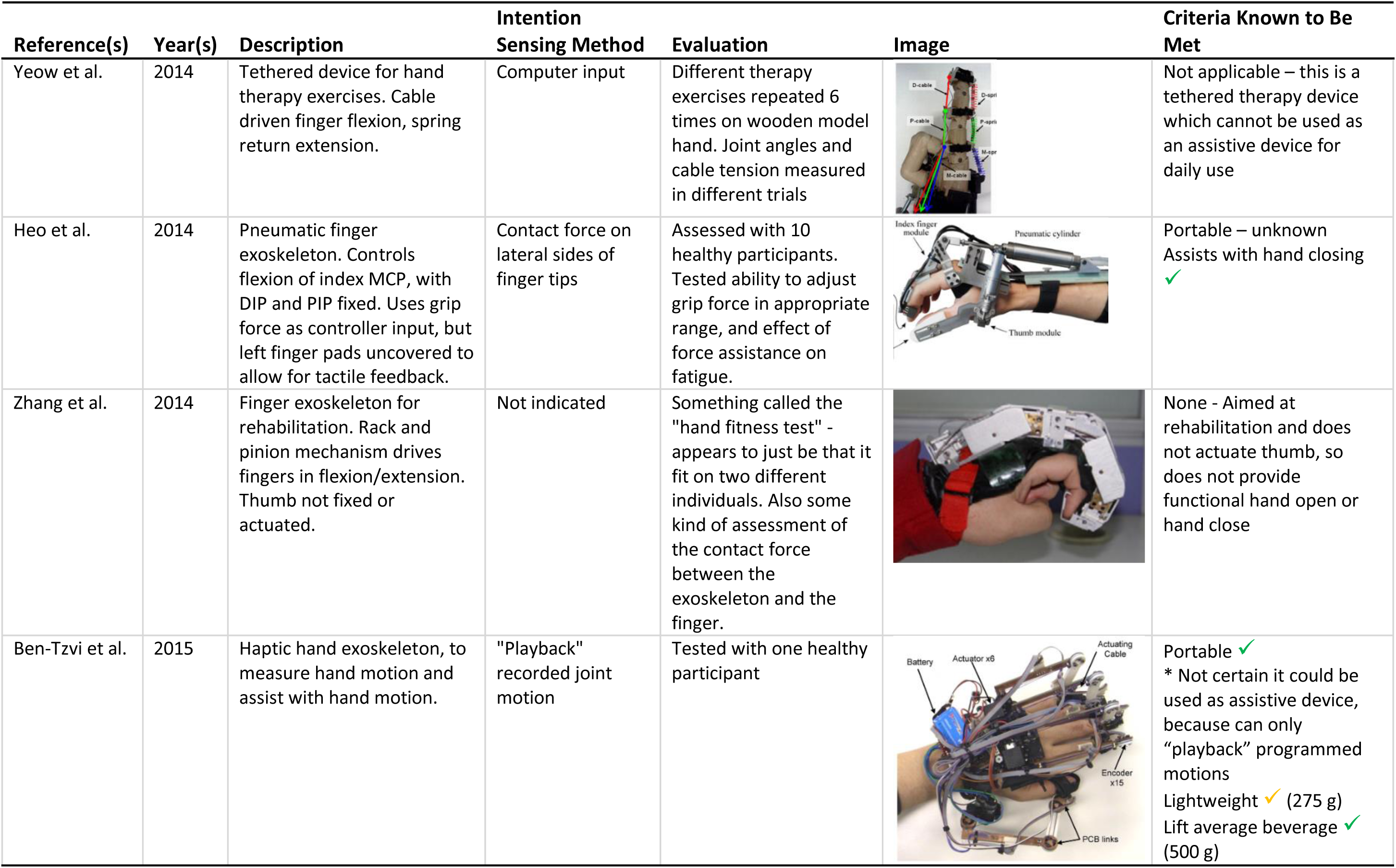

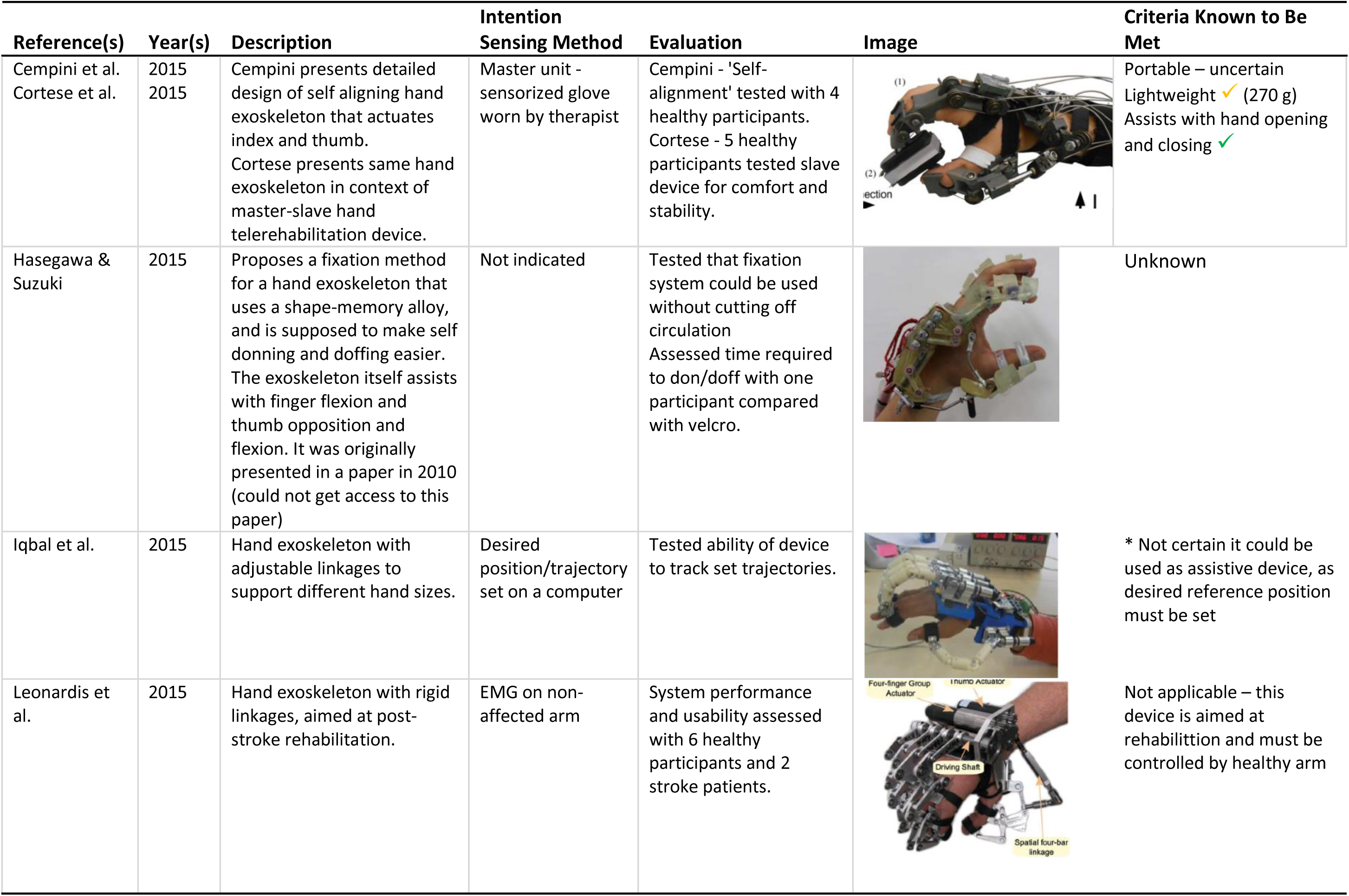

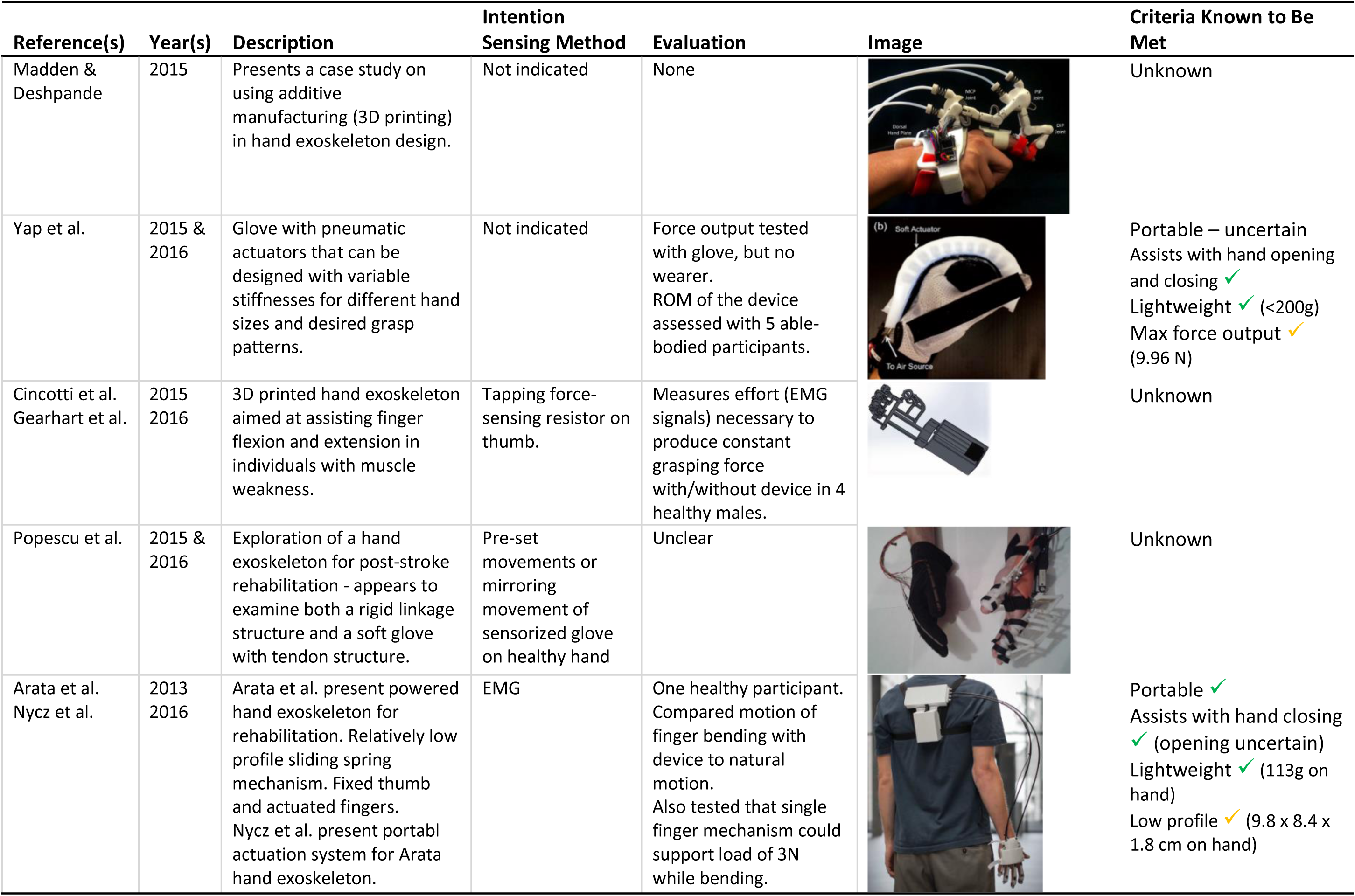

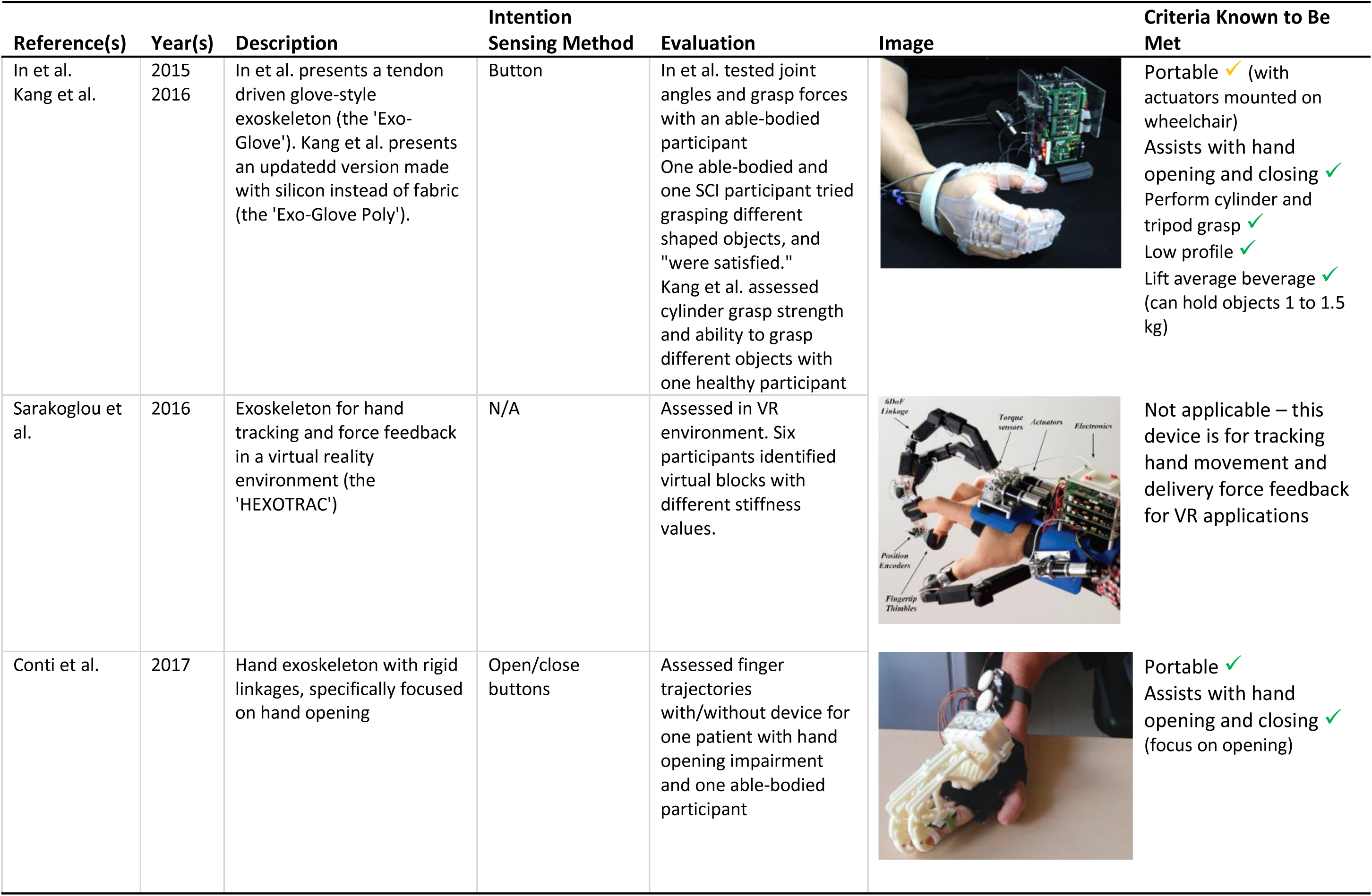

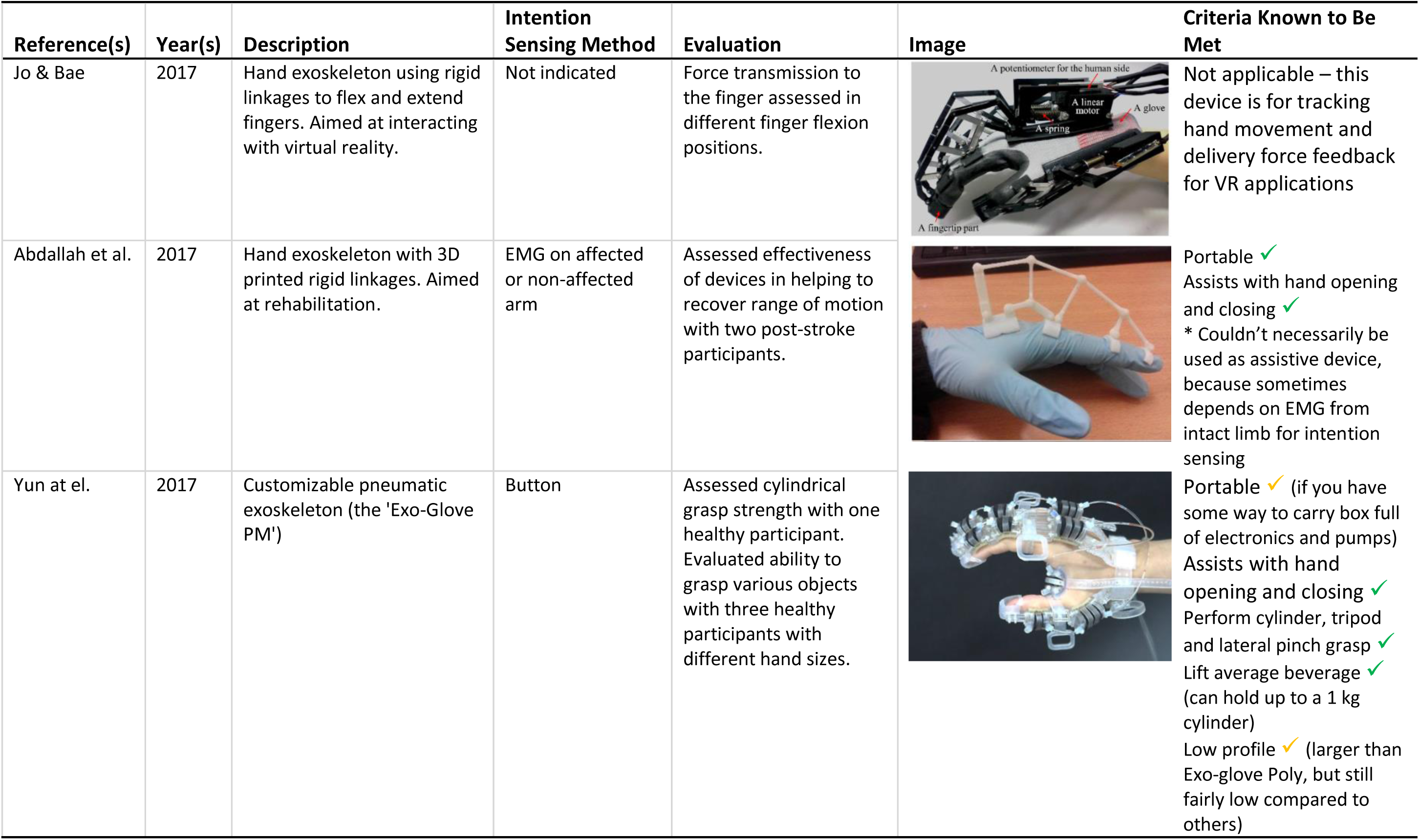

| Reference(s) | Year(s) | Description | Intention Sensing Method | Evaluation | Image | Criteria Known to Be Met |
| --- | --- | --- | --- | --- | --- | --- |
| Jo & Bae | 2017 | Hand exoskeleton using rigid linkages to flex and extend fingers. Aimed at interacting with virtual reality. | Not indicated | Force transmission to the finger assessed in different finger flexion positions. |  | Not applicable – this device is for tracking hand movement and delivery force feedback for VR applications |
| Abdallah et al. | 2017 | Hand exoskeleton with 3D printed rigid linkages. Aimed at rehabilitation. | EMG on affected or non-affected arm | Assessed effectiveness of devices in helping to recover range of motion with two post-stroke participants. |  | Portable ✓<br>Assists with hand opening and closing ✓<br>* Couldn't necessarily be used as assistive device, because sometimes depends on EMG from intact limb for intention sensing |
| Yun et al. | 2017 | Customizable pneumatic exoskeleton (the 'Exo-Glove PM') | Button | Assessed cylindrical grasp strength with one healthy participant. Evaluated ability to grasp various objects with three healthy participants with different hand sizes. |  | Portable ✓ (if you have some way to carry box full of electronics and pumps)<br>Assists with hand opening and closing ✓<br>Perform cylinder, tripod and lateral pinch grasp ✓<br>Lift average beverage ✓ (can hold up to a 1 kg cylinder)<br>Low profile ✓ (larger than Exo-glove Poly, but still fairly low compared to others) |

